# Missing by design: why genomic metadata must be captured at submission, not recovered from the literature

**DOI:** 10.64898/2026.11.03.763936

**Authors:** Soomin Lee, Disha Lodha, Jorge Alvarez-Jarreta

## Abstract

Plant and metazoan genome assemblies are being deposited in the International Nucleotide Sequence Database Collaboration (INSDC) archives at an accelerating rate, but the biological context needed to interpret them, such as ploidy, cultivar, and subgenome structure, is often missing from the record: these attributes are optional in most submission checklists and, when supplied, are entered as free text. We describe the gap in three layers. Ploidy varies within a species and cannot be inferred from taxonomy; chromosome-naming conventions differ between assemblies of the same crop; and that inconsistency is concentrated in species named by subgenome and is not associated with the size of the research community, which points to the lack of a shared convention. As a case study, we built a pipeline that recovers these attributes from the publications linked to an assembly. On a benchmark of 82 assemblies, its current version reported one wrong ploidy, and across 1,059 plant assemblies an earlier, less strict version found the ploidy in a publication for about half; for nearly a third, it found a publication but no statement of ploidy in it. Recovery from the literature can annotate assemblies already deposited but cannot stop new ones from arriving incomplete. We therefore propose that ploidy, cultivar, and subgenome structure be captured at submission as mandatory, controlled-vocabulary attributes, building on existing INSDC and Genomic Standards Consortium checklists.

## 1. Introduction

Plant and metazoan genomes are being deposited in the International Nucleotide Sequence Database Collaboration (INSDC) archives at an accelerating rate, while key biological metadata, such as ploidy, sex or cultivar are not fully captured in a standardised form at submission. Resources such as Ensembl are genome browsers that provide the research community with reference genomes enriched by functional annotation, comparative genomics, and analysis tools [1]. They do not generate assemblies themselves; instead, they draw their sequence data from the INSDC archives [2]. To be reliably interpreted downstream, an assembly’s sequence must be accompanied by biological context such as ploidy, strain or cultivar, sex, and the expected number of chromosomes.

Metadata omissions have direct analytical consequences. Ploidy in particular governs how a genome is computationally handled. Polyploid organisms pose distinct challenges for read mapping and variant calling [3]. The high sequence similarity among subgenomes causes reads to misalign, producing erroneous variant calls [4]. In addition, allelic dosage, the number of copies of each allele, becomes increasingly difficult to resolve as ploidy rises [5]. Chromosome number, likewise, provides a benchmark for assessing an assembly’s completeness and scaffolding. When ploidy or chromosome structure is absent or misattributed, the resulting errors are not overt: a variant caller that assumes diploidy, for example, reports differences between subgenomes as heterozygous sites, yielding output that is plausible yet incorrect. These attributes are prerequisites for comparative genomics rather than descriptive additions, the missing values must currently be recovered through manual curation, in which the associated publications are consulted. A polyploid’s subgenome structure determines whether its duplicated genes are classified as homologues or paralogues. As submission volumes rise, retrieving metadata through manual steps has become a bottleneck.

The metadata problem has two distinct aspects. In many records the information is absent because these attributes are not mandatory in the submission schema. When present, they are usually entered as free text without a controlled vocabulary, so values are inconsistent across records. The responsibility is shared between schema design and submission practice: the fields were made optional because a submitter may not have the information, but they are also left blank when the information is available, either because completing them brings the submitter little direct benefit or because the person making the submission was not given the full details. In many others the information is present in the literature but expressed using conventions that a program cannot reliably parse; these conventions vary from crop to crop and are rarely stated in a form that can be resolved without specialized knowledge of the relevant research community. To analyze the problem systematically, we divide it into three categories. The first is ploidy that varies within a single species. The second is the divergence of chromosome-naming conventions across research communities. The third is a broader pattern that helps explain the second: naming is consistent where a research community has agreed on a shared convention and inconsistent where it has not, largely independent of the community’s size. The second category is harder to resolve by automated extraction than the first, and the third explains why the second arises.

A gap in metadata motivates a position with two components. The absence of metadata from records already deposited must be recovered after deposition. This can be done by manual curation or by asking submitters to update their records, but both depend on individual effort and do not scale; automated extraction from the published literature offers a systematic and scalable route. We present, as a case study, a pipeline built for this purpose and evaluate whether the missing attributes can in fact be recovered at scale. Recovery, however effective, treats the symptom rather than the cause. We therefore argue that it must be paired with intervention at the point of submission: the attributes that curation now labours to reconstruct, namely ploidy, cultivar, and subgenome structure, should be captured at deposition as controlled-vocabulary fields.

## 2. The structural nature of the metadata gap

The outright absence of metadata from the archive has an effect on both sides of the submission process: schemas that leave key attributes optional, and submitters who leave them blank even when the information is at hand. Three features of that process combine to produce the gap.

The first is the design of the submission schema itself. In the INSDC archives, the only attribute of the organism itself that is required for every sample is its taxonomic identity: a sample must be assigned a species-rank name from the NCBI Taxonomy database before it can be registered [6]. Beyond this, the attributes a submitter is required to provide are not fixed but depend on the checklist, or package, chosen at submission time; each checklist specifies its own set of mandatory, recommended, and optional attributes. These sets differ widely from one package to another, and some impose almost no requirements [7]. Biologically important attributes such as ploidy, cultivar, and subgenome structure are either optional or absent and cultivar is rarely required on its own. An assembly can therefore be deposited, and receive an accession, while carrying none of the ploidy, cultivar, or subgenome information that downstream processing may depend on.

The second feature is that the value of these optional fields is not visible to the submitter.

The values themselves are usually known to the submitting group and are often stated with precision in the Materials and Methods section of the accompanying publication, where they are needed to justify the experimental design. But re-entering them into a submission form is a separate, effortful task that yields little direct benefit to the submitter, whose immediate objective is to obtain an accession so they can proceed with the publication. Optional fields, by their nature, are the first to be left blank when effort is scarce. The submission is also not always made by the researchers who generated the data, and the person completing it may not have been given the full details.

The third feature concerns the inconsistency in the structure when it is supplied. Where a field exists and is filled, its contents are frequently entered as free text rather than drawn from a controlled vocabulary. A ploidy value might appear as “tetraploid”, “4x”, or “2n=4x=28”, or be folded into a longer descriptive phrase; a cultivar name might be abbreviated, spelled inconsistently, or replaced by an internal code. Free-text entry admits synonyms, formatting variation, and ambiguity that a downstream program cannot reliably normalize, so that even metadata that is present may not be usable without human interpretation. It is here that the first aspect of the problem, absence, begins to shade into the second, the resistance of recorded information to automated interpretation, which the following sections examine in detail.

## 3. Layer one: ploidy varies within a species

The most tractable category arises because ploidy is not a fixed property of a species. One approach to a missing ploidy value would be to look it up in a reference resource indexed by species name, such as Genomes on a Tree (GoaT) [8]. This is adequate for species whose ploidy is uniform, but unreliable for cultivated species in which ploidy varies between cultivars because a species-level value cannot distinguish them. Apple is a clear example: most cultivars of *Malus domestica* are diploid (2n = 2x = 34), yet several commercially important ones, such as ‘Bramley’s Seedling’ and ‘Jonagold’, are triploid (2n = 3x = 51) [9]. The species name does not distinguish them; only the cultivar does. The same holds across crops: cultivated potato includes autotetraploid varieties alongside diploid landraces of *Solanum tuberosum* [10], and banana spans diploid, triploid, and tetraploid cultivars within the *Musa* genus [11]. In each case, ploidy follows the specific cultivar or breeding line rather than the taxon. In our benchmark (Section 6), GoaT species-level values were correct for 30 of the 31 assemblies to which they were applied; the exception was a *Saccharum spontaneum* line (AP85-441) that is tetraploid, whereas GoaT records the species as octoploid. Identifiers broader than the species fail for the same reason: the common name “wheat” covers both hexaploid bread wheat, *Triticum aestivum* (2n = 6x = 42), and tetraploid durum wheat, *Triticum turgidum* subsp. *durum* (2n = 4x = 28) [12,13], so a lookup keyed on a common name or genus would confuse the two.

Because ploidy is a property of the sequenced individual rather than its species, it cannot in general be inferred from taxonomy; it must be read from that individual’s record, typically the assembly’s own publication. This is the main reason the value must be recovered from the literature rather than a reference table. Ploidy can also be estimated directly from sequencing reads with k-mer based methods such as GenomeScope 2.0 and Smudgeplot [14], but running them on the raw reads of every assembly is costly, their estimates are less certain for allopolyploid and highly heterozygous genomes, and the results are not recorded in the archive either. This category is also the most extractable of the three, because once found in the text the value is typically stated explicitly and does not depend on community-specific convention.

## 4. Layer two: naming conventions fragment across communities

In the second category, the metadata is present in the literature but written in conventions that a program cannot parse without specialized knowledge, and that are not applied uniformly from one assembly to the next. This is most evident in chromosome naming. In bread wheat, a chromosome is named by its homoeologous group (1 to 7) and its subgenome (A, B, or D), giving labels from 1A to 7D, with sequence that cannot be placed collected under a label such as “ChrUn” [12]. Thus, the string “1A” is meaningful only to a reader who is already familiar with this scheme. However, these conventions are not consistent even within a single crop. In peanut, chromosomes are numbered 1 to 20 in some assemblies, labelled A01 to A10 and B01 to B10 by subgenome in others, and prefixed as Arahy.01 to Arahy.20 in yet others, so the same physical chromosome carries different names across assemblies [15–17]. *Brassica napus* follows yet another pattern, with chromosomes labelled A01 to A10 and C01 to C09 after its two progenitor genomes, while legacy linkage-group names (N1 to N19) persist in older records [18]. Because no single convention applies, and each requires the knowledge of the group that established it, a program cannot resolve chromosome identifiers by a general rule; it must maintain a separate mapping for each crop and, in some crops, for each assembly.

## 5. Layer three: shared conventions, not community size, determine standardization

Whether chromosome names are standardized appears to depend on whether a community has agreed on a shared convention. We examined the chromosome names of 1,005 chromosome-level GenBank assemblies representing 111 plant species, each with at least five chromosome-level assemblies in GenBank. Naming was consistent within most species: for 86 of the 111, every sampled assembly used the same naming structure (the sequence of numbers and single letters in each name, ignoring word prefixes such as “Chr” or “LG”, separators, and zero padding), and consistency was not associated with community size, measured as the number of PubMed records for the species (Spearman ρ = −0.06, permutation p = 0.56, n = 111). By exact format, which also distinguishes prefixes and zero padding, 76 of the 111 species were consistent, and this measure was not associated with community size either (ρ = −0.16, p = 0.10). Inconsistency was instead concentrated in species whose chromosomes are named by subgenome, such as polyploids and their close relatives. Bread wheat, for which the International Wheat Genome Sequencing Consortium (IWGSC) established a reference assembly and the 1A to 7D nomenclature [12,19], used one naming format in all 12 assemblies sampled. *Brassica napus*, whose research community is also large (6,777 PubMed records, against 12,882 for wheat), used three formats across 10 assemblies (A1, A01, and 1 to 19), and peanut used three (1 to 20, A01 to B10, and Arahy.01 to Arahy.20) [15–18,20]. The inconsistency that frustrates automated extraction therefore arises where subgenome structure calls for a naming convention and none has been agreed. Wheat shows that a polyploid can be named consistently once a convention exists, and *B. napus* shows that a large community does not produce one by itself.

Because what is missing is agreement, the inconsistency is open to a coordinated remedy rather than to ever more elaborate extraction.

## 6. Recoverability as evidence: a case study

If the missing attributes are, as argued, present in the literature but unevenly captured, then it should be possible to recover them by reading the literature directly. We built a pipeline as shown in Figure 1 that, given an assembly accession, retrieves publications linked to the assembly, reads its text together with any supplementary files, and extracts ploidy, chromosome number, cultivar or strain, and sex. The pipeline is designed to favour precision over coverage: it grounds every value in a quoted passage, combines a rule-based extractor with a keyword and dense-retrieval ensemble and an optional language-model layer, discards papers that are matched only by species name, withholds values whose evidence is weak or conflicting, and reports no value when the text provides no evidence, rather than inferring one from prior expectation. These safeguards were introduced in stages, and the results below refer to three versions of the pipeline. The first version still accepted papers matched only by species name, including papers that mention the species only in their full text. The second accepted only papers tied to the assembly, as in Figure 1. The current version adds the precision guard: unless a ploidy is backed by a chromosome formula consistent with the assembly, it is withheld when its evidence is weak or conflicting, when the statement does not refer to the sequenced organism, or when it contradicts GoaT’s directly curated value for the species. Each step traded coverage for precision.

**Figure 1.**
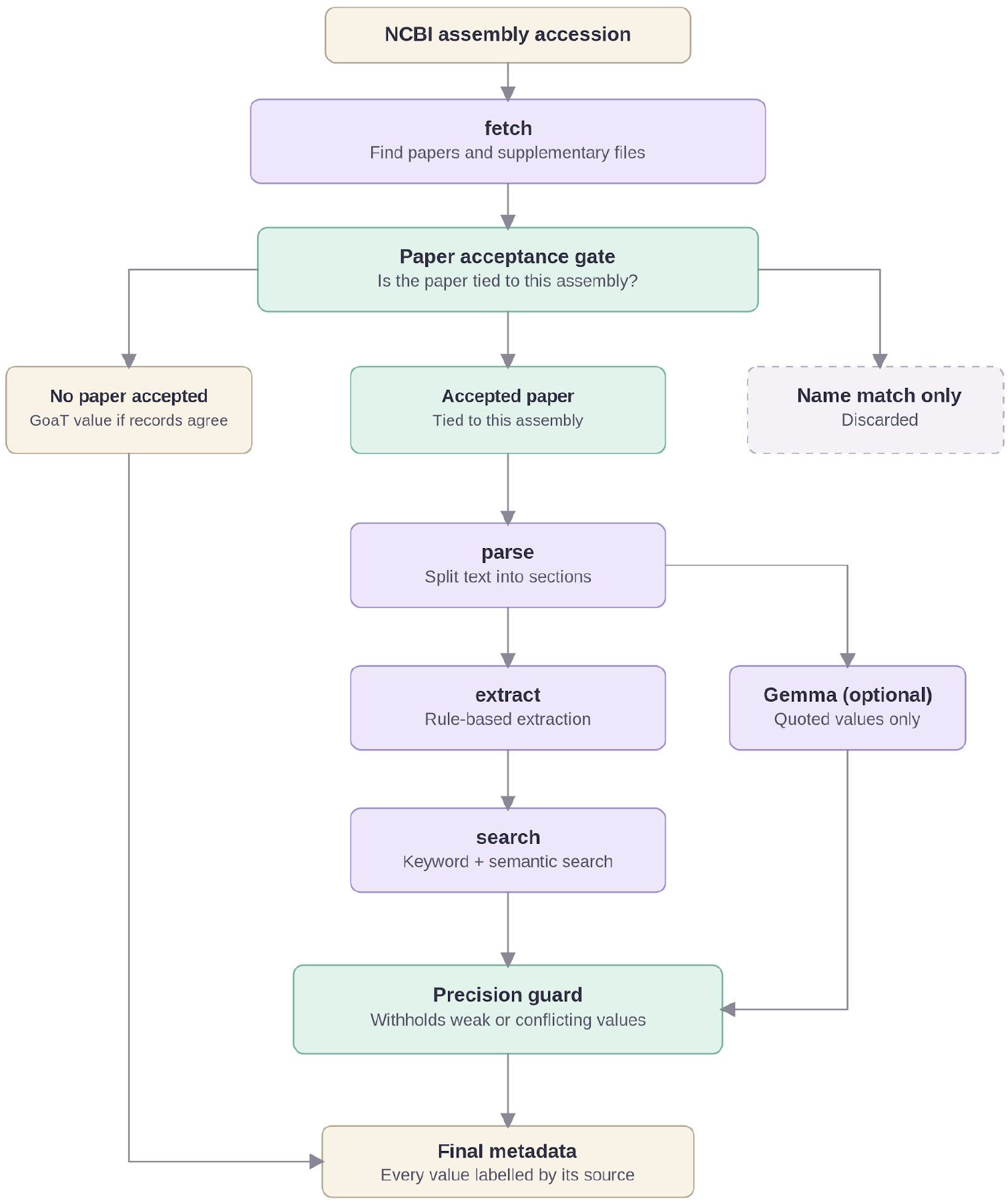
Overview of the literature-based metadata extraction pipeline. A paper is accepted only if NCBI links it to the assembly’s accession, if it is the BioProject reference paper and names the organism, or if the accession appears in its text; papers matched only by species name are discarded. Accepted text is processed by rule-based extraction, keyword and semantic search, and an optional Gemma model, and a final check withholds values whose evidence is weak or conflicting. When no paper is accepted, the GoaT species-level ploidy is reported as a reference value, provided GoaT’s direct record for the species agree.

We evaluated the pipeline in two ways. First, against ploidy established by manual curation. On 11 curated plant assemblies, the pipeline recovered the curated value for 10 using rule-based extraction and vector search alone. To test precision directly, we assembled 82 assemblies, 37 of them edge cases chosen to provoke false positives, including diploid relatives of polyploids, doubled-monoploid lines, haploid fungi and high-ploidy crops, and checked each label against the assembly’s organism in NCBI and the published literature. On this set, the current version reported a wrong ploidy for 1 assembly, compared with 14 for the second version, at the cost of leaving more values empty (38 compared with 22). Of the 44 values the current version reported, 43 were correct; 30 of these came from GoaT and 13 from a publication. Second, to test behaviour at scale, we ran the first version of the pipeline on the first 1,059 chromosome-level plant assemblies returned by NCBI Datasets, covering 1,057 species, without further selection. Every assembly returned a record. A ploidy was read from a publication for 512 assemblies (48%) and taken from GoaT for a further 201 (19%), spanning haploid to decaploid. Because the first version also accepted papers matched only by species name, these figures are an upper bound on what the current version would recover. With no ground truth at this scale, this run measures coverage and robustness, not accuracy.

The assemblies left without a value are themselves informative. Of the remaining 346, only 12 had no publication that could be found; the other 334 had a retrievable publication in which the pipeline found no statement of ploidy. For about half of the assemblies, then, the ploidy the archive lacks was present in the literature, while for nearly a third the pipeline found a publication but no statement of ploidy in it. Nor do the assemblies themselves hint at the missing value. Of the 346, 330 name their chromosomes with plain numbers or linkage groups (such as 1 or LG01) and 4 add a single letter to each number (such as C1), none of which says anything about ploidy. Only 12 use labels that separate two or more copies of each chromosome (such as 1A and 1B); these point to a polyploid or to a haplotype-resolved hybrid without saying which, although three of them separate three or six copies. The gap is therefore not only one of capture from the literature into the archive: for many assemblies the value may never have been written down in a retrievable form, which is the strongest argument for recording it at submission.

Recovery from the literature is necessary, but it is not enough on its own. It works only after an assembly is deposited, and only when a publication exists and can be found. The hardest cases show the limit. For *Agapanthus africanus*, the correct tetraploid value (2n = 4x = 30) appears only in the full text of a methods paper on genome size estimation [21]; we could find no genome paper for the species, and the assembly’s archive record links no publication. The first version recovered it through its full-text search, but the later versions reject that paper, because nothing ties it to the assembly, and report no value. The trade-off is deliberate: the same broad search that found the *Agapanthus* value also produced false positives, such as an *Arabidopsis thaliana* assembly reported as tetraploid from a paper on *Dioscorea* [22]. Such cases, along with publications that are paywalled or that do not exist, set a limit on how much any after-the-fact method can achieve. Recovery can therefore clear the backlog of assemblies already in the archive, but it cannot prevent the next assembly from arriving without its metadata. Preventing that requires acting earlier, at the point of submission. In total, the pipeline was benchmarked on 11 manually curated plant assemblies and on 82 edge-case assemblies, and run at scale on 1,059 assemblies from 1,057 plant species: every run completed without failure, taking a median of about two minutes and under 2 GB of memory per assembly, and the full set took 6.6 hours on a single server processing five to six assemblies in parallel.

## 7. A path forward: capturing metadata at submission

Metadata should be captured at deposition, as mandatory attributes drawn from standardized vocabularies, rather than reconstructed afterwards. This reverses each of the three features described in Section 2: permissive schemas, weak incentives, and free-text entry. Ploidy, cultivar, and subgenome structure would move from optional to required in the submission checklists that apply to plant and metazoan assemblies. Their values would be recorded as standardized terms rather than free text, so that “tetraploid”, “4x”, and “2n = 4x = 28” resolve to a single value. Structured capture at submission also collapses the three layers that make recovery hard. Ploidy that varies within a species is recorded for the deposited individual, not inferred from its taxon. The chromosome-naming schemes that differ between assemblies are irrelevant to a standardized ploidy or subgenome attribute. For chromosome-level use, the subgenome of each chromosome would also need to be recorded, for example as a field in the assembly’s chromosome list, so that software can match chromosomes across assemblies without parsing their names. And uneven standardization across communities no longer governs data quality, because the checklist enforces one structure on every submission.

The archives already make attributes mandatory once their value for downstream reuse is widely accepted. INSDC sample checklists make collection date and geographic location mandatory [23], and the Genomic Standards Consortium’s MIxS checklists provide an established framework of required, recommended, and optional attributes that already defines a ploidy term for eukaryotic genomes and into which cultivar and subgenome structure would fit [24]. The mechanism and the standards therefore already exist; extending them to these attributes is a question of policy, not of technical feasibility. The cost to the submitter is minimal, since the submitting group knows these values at deposition, usually before the paper is published. When a value is not known, the missing-value terms that INSDC already accepts for collection date and location, such as “missing: not collected”, let the submitter say so explicitly [23]. Extraction from the literature can clear the assemblies already deposited, but only capture at submission prevents the backlog from re-forming.

## 8. Conclusion

Ploidy, cultivar, and subgenome structure, the context required for correct variant interpretation and comparative analysis, are often absent from the archive records of plant assemblies. For ploidy, the literature fills only part of this gap: in our case study, a publication supplied the value for about half of the assemblies, while for nearly a third the pipeline found a publication but no statement of ploidy in it. This gap is structured: ploidy varies within a species and cannot be inferred from taxonomy; chromosome-naming conventions differ between assemblies and resist a general parser; and that inconsistency arises where no shared convention has been agreed, marking it as a problem of coordination rather than a technical limitation.

A literature-extraction pipeline shows that these values can be recovered at scale where a publication states them, but such recovery is inherently retrospective: it can annotate assemblies already deposited, but cannot prevent new submissions from arriving incomplete. A durable solution requires capturing ploidy, cultivar, and subgenome structure as mandatory, standardized attributes at the point of submission, for which INSDC checklists provide an established precedent.

Two lines of work follow. First, controlled vocabularies for these attributes should be defined and integrated into existing submission checklists, in coordination with the sequence archives and the Genomic Standards Consortium, and piloted on the polyploid and variable-ploidy crops where these attributes are most often misread, such as those examined here. Second, extraction should be applied to the existing backlog, with recovered values returned to curators as evidence-linked suggestions rather than assertions. Together, retrospective recovery and prospective capture would convert a recurring curation burden into a problem solved once, at the point where the information originates.

## Data availability

The genomic metadata and assembly information analyzed in this study were obtained from publicly available resources, including the NCBI Assembly database and Genomes on a Tree (GoaT). Representative examples of ploidy and chromosome-naming conventions are summarized in Table 1. The literature-based metadata extraction pipeline is publicly available as described in the Code availability section.

**Table 1.**
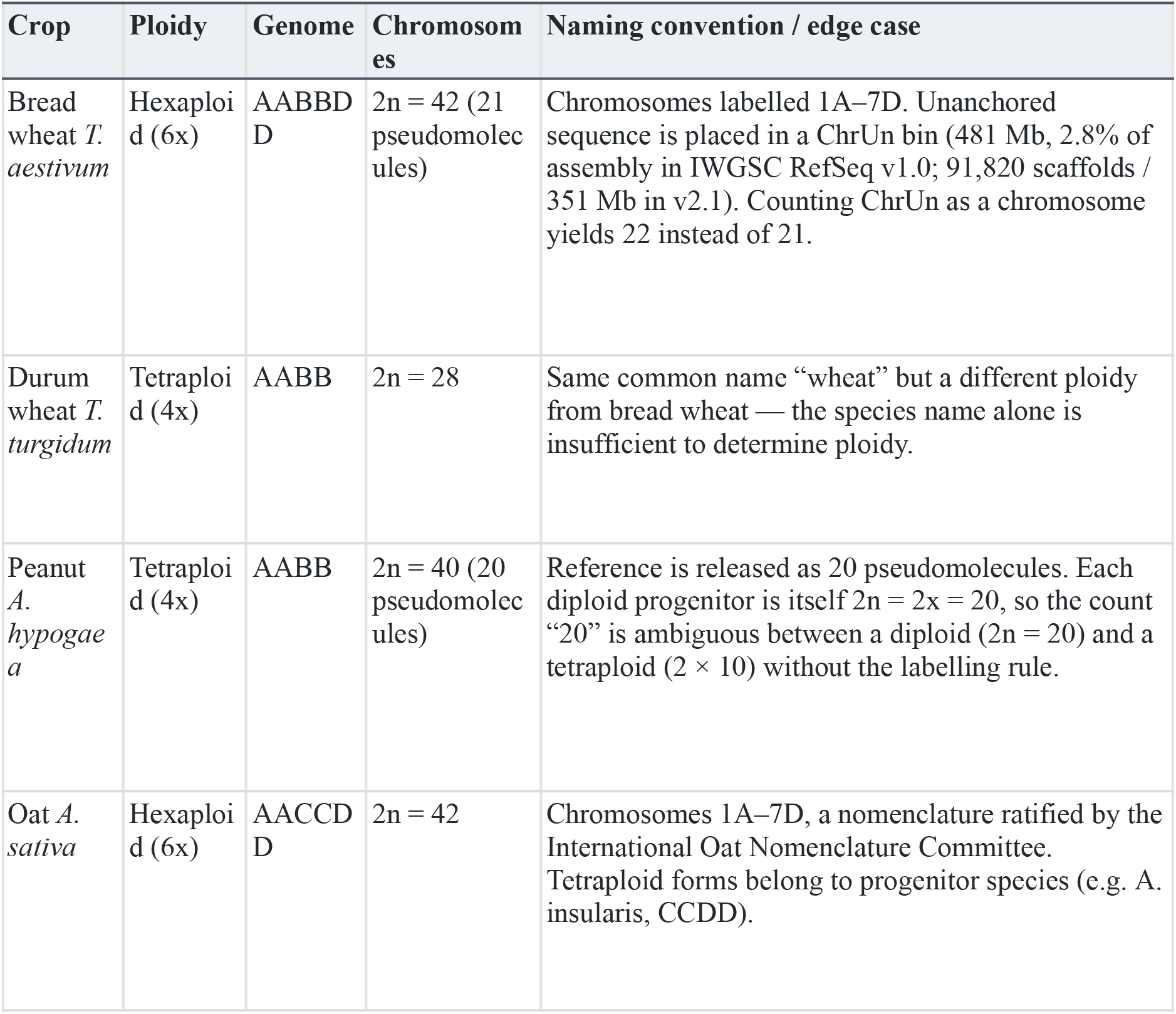
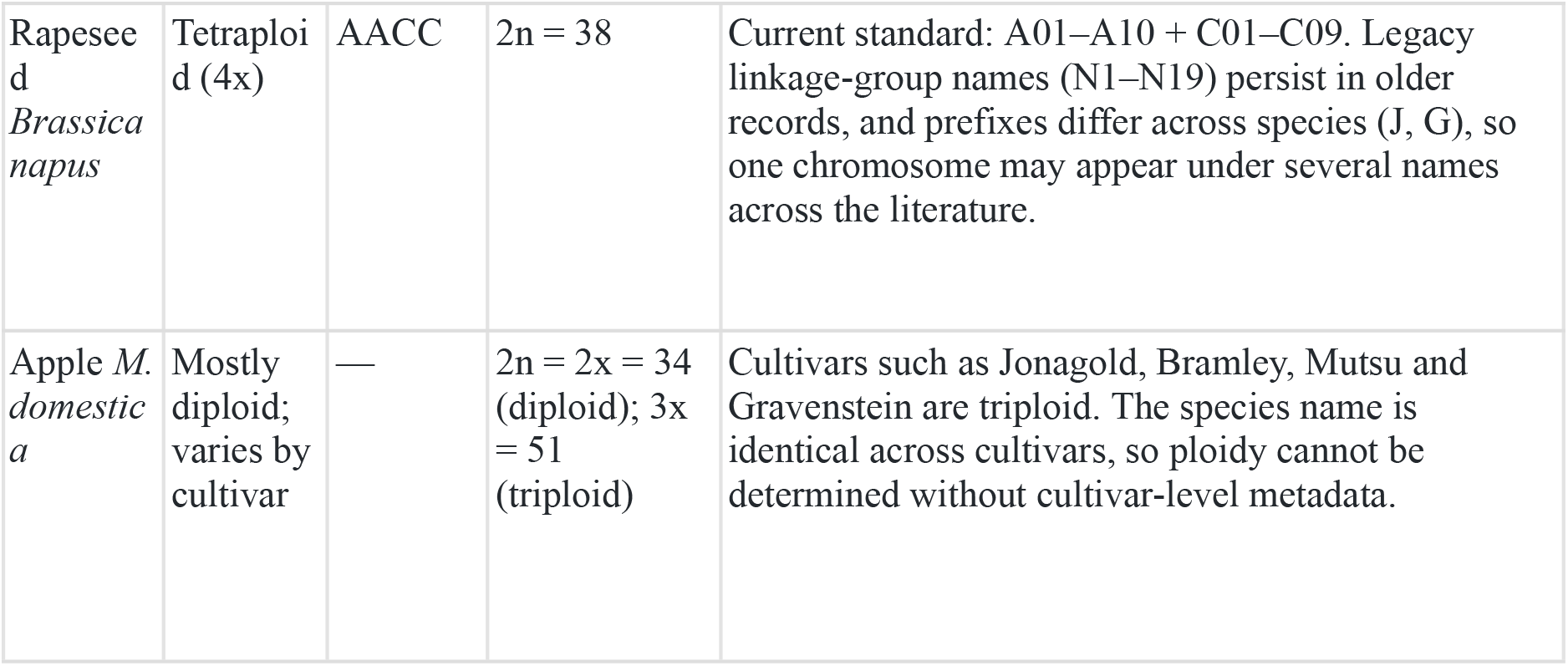
Ploidy and chromosome-naming conventions across representative polyploid and variable-ploidy crops, illustrating why species name and chromosome count alone are insufficient for automated metadata inference.

| Crop | Ploidy | Genome | Chromosomes | Naming convention / edge case |
|---|---|---|---|---|
| Bread wheat *T. aestivum* | Hexaploid (6x) | AABBDD | 2n = 42 (21 pseudomolecules) | Chromosomes labelled 1A–7D. Unanchored sequence is placed in a ChrUn bin (481 Mb, 2.8% of assembly in IWGSC RefSeq v1.0; 91,820 scaffolds / 351 Mb in v2.1). Counting ChrUn as a chromosome yields 22 instead of 21. |
| Durum wheat *T. turgidum* | Tetraploid (4x) | AABB | 2n = 28 | Same common name “wheat” but a different ploidy from bread wheat — the species name alone is insufficient to determine ploidy. |
| Peanut *A. hypogaea* | Tetraploid (4x) | AABB | 2n = 40 (20 pseudomolecules) | Reference is released as 20 pseudomolecules. Each diploid progenitor is itself 2n = 2x = 20, so the count “20” is ambiguous between a diploid (2n = 20) and a tetraploid (2 × 10) without the labelling rule. |
| Oat *A. sativa* | Hexaploid (6x) | AACCDD | 2n = 42 | Chromosomes 1A–7D, a nomenclature ratified by the International Oat Nomenclature Committee. Tetraploid forms belong to progenitor species (e.g. A. insularis, CCDD). |
| Rapeseed *Brassica napus* | Tetraploid (4x) | AACC | 2n = 38 | Current standard: A01–A10 + C01–C09. Legacy linkage-group names (N1–N19) persist in older records, and prefixes differ across species (J, G), so one chromosome may appear under several names across the literature. |
| Apple *M. domestica* | Mostly diploid; varies by cultivar | — | 2n = 2x = 34 (diploid); 3x = 51 (triploid) | Cultivars such as Jonagold, Bramley, Mutsu and Gravenstein are triploid. The species name is identical across cultivars, so ploidy cannot be determined without cultivar-level metadata. |

## Code availability

The pipeline is implemented in the ensembl.io.genomio.literature module of a fork of the Ensembl ensembl-genomio package (https://github.com/soominhildelee/ensembl-genomio, branch literature-post-gsoc-refinements).

## Author contributions

S.L. developed the pipeline, performed the analyses, and wrote the first draft. D.L. and J.A. supervised the work and revised the manuscript. All authors approved the final version.

